# “A compressed representation of spatial distance in the rodent hippocampus’’

**DOI:** 10.1101/2021.02.15.431306

**Authors:** Daniel J. Sheehan, Stephen Charczynski, Blake A. Fordyce, Michael E. Hasselmo, Marc W. Howard

**Affiliations:** Boston University

## Abstract

Principal cells in the rodent hippocampus often fire in response to traversal through a specific spatial location (place cells), as well as elapsed time during an imposed temporal delay or after stimulus offset (time cells). Sequences of time cells unfold rapidly at first, with many time cells with narrow time fields. As the triggering event recedes into the past, time cells are fewer and have broader fields. This means that the representation of time in the hippocampus is compressed with greater resolution for time points near the present. Using tetrode recordings we measured individual CA1 units while rats traveled along a track that could be changed in length. Consistent with previous results, most place cells coded for distance from the starting point of the trajectory. Critically, place cells became less numerous and showed gradually widening fields with distance from the starting location. These results suggest that as the animal leaves a landmark, the hippocampal place code forms a compressed representation of distance from the starting location. The representation of time and space in the hippocampus have similar properties suggesting that they arise from similar computational mechanisms.

**Significance Statement:** The hippocampus represents relationships between events in time and space. It has been hypothesized that temporal and spatial relationships are the result of a common computational mechanism. Previous work has shown that the representation of time in the hippocampus is compressed, with less neural resolution for more temporally remote events, consistent with the observation that temporal memory is worse for events further in the past. This paper shows an analogous result for spatial relationships. Place cells coded for distance from the start of a journey. As distance increased, place fields became broader and less numerous, showing a decrease in spatial resolution. This result suggests a unified coding scheme for the dimensions of time and space in the rodent hippocampus.

## Introduction

The hippocampus is critical for episodic memory (Scoville & Milner, 1957) as well as the relational structure for memories of personal experiences (Eichenbaum, 2017a). Overlapping populations of pyramidal cells in dorsal CA1 are believed to represent position during physical traversal of space via place cells (O’Keefe & Doskovsky, 1971), as well as elapsed time during a delay via time cells (Pastalkova et al., 2008; MacDonald et al., 2011) providing a neural basis for the organization of experience as a function of spatial and temporal relationships. It has been proposed (e.g., Buzsaki and Tingley, 2018; Eichenbaum, 2017a, Hasselmo, 2012) that representations of space and time in the hippocampus reflect a common computational mechanism, with similar properties.

Time cell sequences recorded from the rodent hippocampus exhibit compression for firing fields that span the delay following an event (Kraus et al., 2013; Mau et al., 2018), meaning that both the number of fields decreases and the field size widens as a function of the delay. These properties have been postulated to be important in accounting for behavioral properties of memory (Howard & Eichenbaum, 2013), such that the decreasing resolution of the time code for events further in the past contributes to forgetting with the passage of time. Compressed time cells have been observed in CA1 while holding the animal fixed in space via stationary running (Kraus et al., 2013; Salz et al., 2016; Robinson et al., 2017; Mau et al., 2018) or head-fixed restraint (Terada et al., 2017, Taxidis, et al., 2020), during a working memory delay.

If the representations of space and time in the hippocampus share a common computational mechanism, then we would expect the place code to also be compressed. Time cells show an increase in field width as a function of temporal distance from the present. In order to ask the analogous question for place cells it is necessary to establish a reference point for the population. Indeed, it has long been appreciated that place cells can code for conjunctions of relationships to distant landmarks (e.g., O’Keefe & Burgess, 1996). Principal cells in the CA1 region have been demonstrated to show spatial anchoring to both static and moveable landmarks (Gothard et al., 1996; Gothard et al., 2001). In experiments on a linear track with a movable start box, the place code can transition rapidly from a reference frame that attaches to the box to a reference frame that is fixed in room coordinates (Redish et al., 2000; Jezek et al., 2011; Kelemen & Fenton 2016; Kay et al., 2020). Previous experimental work has shown potential examples of non-uniform place field allocation along a linear track apparatus (Gothard, et al., 1996; 2001; Bjerknes et al., 2018), yet there has been little systematic testing of the distribution of place fields in a 1-dimensional environment. Recently, the spatial tuning of CA3 place fields of gerbils in a virtual environment showed sensitivity to modulations of visual cue gain relative to proprioceptive cues, resulting in place field widening and density of fields nearest trajectory ends, along the length of the VR track (Haas et al., 2019). However, whether the coding of space in region CA1 of the hippocampus has compression-like tuning profiles, as has been observed for time and hypothesized for both space and time (Howard & Eichenbaum, 2013), has not yet been directly tested.

In this present study, the spatial coding characteristics of principal CA1 neurons in the hippocampus of freely navigating rats is investigated in a one-dimensional (1D) environment in the same manner as has been done for temporal intervals. During goal-oriented linear track running the neural dynamics at both the single unit and population level are compared in relation to different spatial reference frames. In sum, this study addresses whether the hippocampus represents the dimensions of time and space in a similar manner, suggesting a unified computational framework for hippocampal coding of experience.

## Materials & Methods

### Subjects

Subjects were four male Long-Evans rats (Charles River) weighing between 350 and 500 grams at the start and duration of the experiment. All animals were single housed and maintained on a 12 h light/dark cycle (lights on 8:00 A.M. to P.M.). Behavioral training and testing were conducted exclusively during the light phase of this housing cycle. All animals had *ad libitum* access to food in their home cage. During behavioral training and testing, rats were placed on water-restriction when behavioral training required it and returned to *ad libitum* water access, along with food, during weekends. Animals were maintained at a minimum (85%) of their *ad libitum* feeding body weight during all behavioral training and testing periods. Procedures were conducted in accordance with the requirements set by the National Institutes of Health and Boston University Institutional Animal Care and Use Committee (IACUC).

### Linear Track environment

The behavioral training and testing environment was a custom-built wooden linear track apparatus (223.53 l x 10.8 w cm) elevated 96.52 cm off the ground as illustrated in Figure 1. One end of the linear track had a fixed reward point (waterport) embedded in a block of wood (8.8 w x 8.8 l x 5 h cm). The opposite end of the linear track consisted of a moveable wooden box (26.67 l x 31.75 w x 27.94 h cm) with a waterport positioned 7.62 cm from the rear of the chamber and a sliding wooden door on the front (track facing) side of the box, to allow for enclosure of the animal during the 10 second Inter-TriaI-Interval (ITI) period. When called for, this start box could be positioned at six equal intervals along the track, each separated by a distance of 27.94 cm. The longest distance from the front edge of the box to the opposite end of track reward point measured 180.34 cm. The entire track and box was painted black, except for two distinct visual cues on the interior side walls of the movable box. Water delivery was controlled by custom MATLAB script that interfaced with a National Instruments DAQ box (model #6501) allowing for water to be dispensed by solenoids connected to a gravity fed water reservoir. The linear track environment was situated in the middle of the recording room and enclosed on one side by movable dark black rubber curtains, allowing for viewing of distal room cues by the animal.

**Figure 1:**
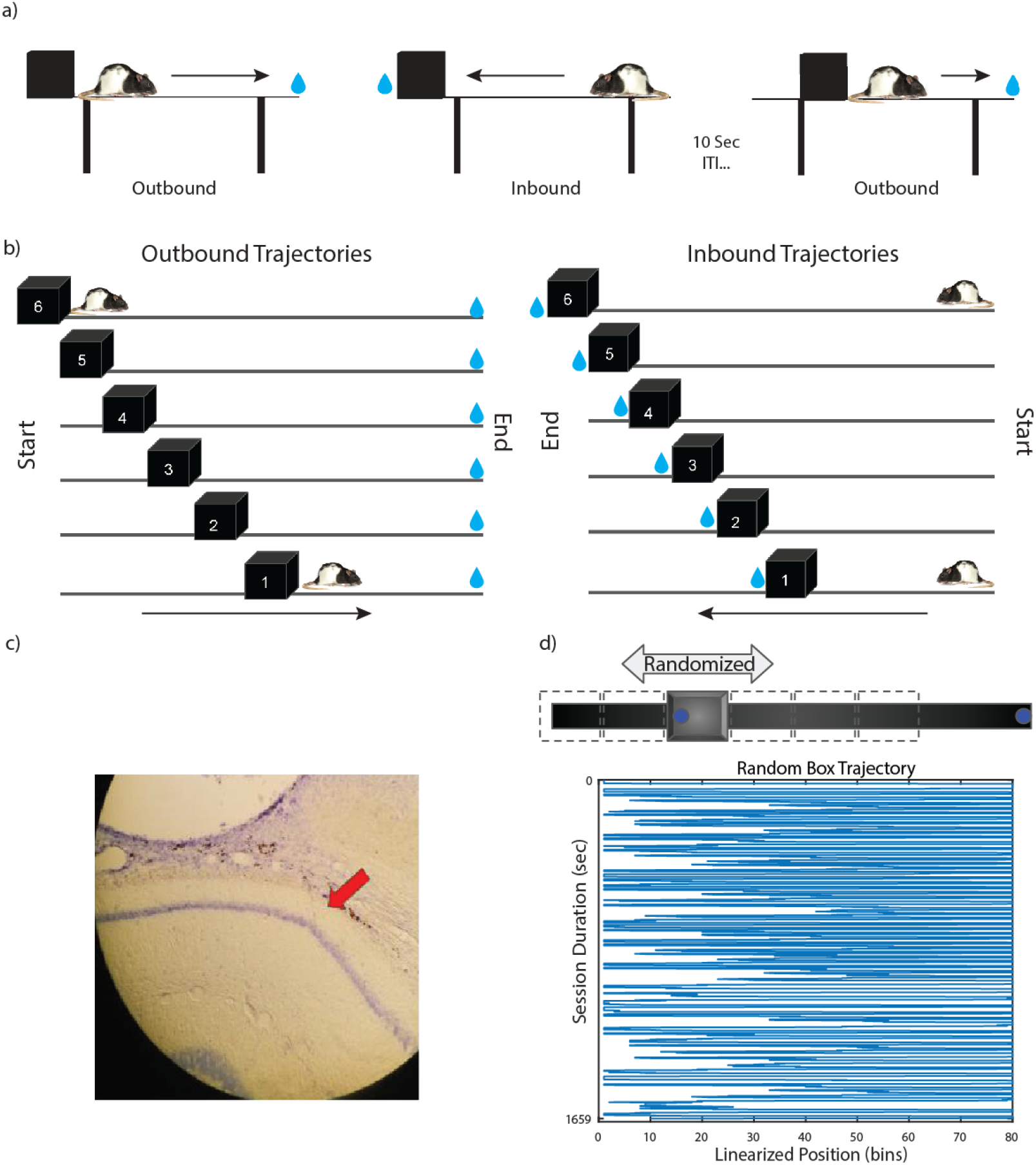
Experimental protocol overview. a) Diagram of linear track apparatus showing reward locations and depicting trajectories during active behavior. Individual trials are separated by a 10 second ITI. Water rewards are dispensed after completion of each directional trajectory. b) Depiction of box positioning along the linear track apparatus for both outbound (left) and inbound (right) trajectories during the randomized start box recording sessions. c) Histological image of dorsal CA1 showing representative locations for neural recordings (red arrow indicates tetrode track). d) Graphic of the randomized box location protocol (above) and an example plot of an animal’s linearized position (below) during a representative random box recording session.

### Linear Track Behavioral Protocols

Subjects were first exposed to and shaped to run along the linear track before surgical implantation. This was achieved by a training protocol that consisted of first placing the animal into the closed “start box” to develop familiarity. The door was slid open, allowing entry onto the track surface, allowing free exploration of the apparatus. Once animals reached the opposite end of the track (track end), a water reward was dispensed. Completing a traversal of the entire track resulted in another water reward being dispensed from a waterport located at the rear of the start box. Training continued until animals demonstrated reliable and consistent track running behavior, by leaving the box coincident with the front door being opened as well as returning back to the initial starting location to receive a second water reward. Once subjects demonstrated appropriate track running behavior they underwent surgical implantation and a subsequent recovery period before being re-exposed to the track environment. A brief reshaping and training period was required, in order for them to demonstrate appropriate track running behavior in line with requirements for neural recording sessions.

For every neural recording session, the start box position always began at the longest track length (*box position 6*). Condition one (*baseline*) consisted of the rats running back and forth along the track, with the box fixed for all trials at the longest track length (*box position 6*), with the room moderately lit, in order for the rat to view both local and distal room cues. Each trial began with the opening of the start box door and the animal was allowed to journey “outbound” along the linear track towards the opposite end of the track, in order to receive a water reward. After the water reward was consumed, the animal self-initiated the return (inbound) journey. Once the rat completed the inbound journey, making contact with the front edge of the start box, a water reward was dispensed, signalling the end of the inbound trajectory. Once the rat fully entered the start box, the front door of the box was slid closed and a 10 second inter-trial-interval commenced. After 10 seconds had elapsed, the start box door was once again opened, allowing for the rat to self-initiate the next trial with an “outbound” trajectory. Each recording session duration for “stable” box sessions was based on the completion of 50 trials or until the rat failed to leave the box after 30 seconds on multiple neighboring trials.

In the second condition the start box was pseudo-randomly shifted between the six possible box locations for each trial with the room moderately lit. In this condition the first trial always started with the start box being placed at the longest track location (box position 6). Similar to the baseline (stable box) condition, the rat was allowed to freely consume the water reward and explore the immediate end segment of the linear track, before turning around to initiate their return journey (inbound trajectory) back into the open start box to receive another water reward. Movement of the box consisted of carefully sliding the box along the track to the next trial’s starting position during the 10 second ITI, with the rat inside the closed start box. The next trial began with the opening of the front door of the start box, allowing the animal to exit the box from the new starting location along the track, to which they would return after completing both the outbound and inbound traversals of the track. Random box recording sessions lasted for the duration of 60 total trials, or until the rat failed to leave the box after 30 seconds on multiple subsequent trials.

In addition to the above behavioral protocols, animals were also exposed to recording sessions that subjected the animals to various lighting conditions. For these recording sessions, lighting was manipulated in an alternating blocked manner, consisting of the lights being either ‘ON’ or ‘OFF’ for a duration of 10 full trials before being switched. Both the stable and moving box conditions were used in tandem with the blocked lighting manipulation for these additional recording sessions. In general, recording sessions with variable lighting occurred after data collection of the lights ‘ON’ version of running the specific behavioral tasks, as well as on separate days. Results and findings from these variable lighting recording sessions will be reported elsewhere.

### Surgical implantation

Anesthesia was induced by inhalation of 5% isoflurane (Webster Veterinary Supply) in oxygen and was maintained at 1.5%-3% throughout the entirety of surgery. Before surgery animals were injected with the analgesic Buprenex (buprenorphine hydrochloride, 0.03 mg/kg i.m.; Reckitt Benckiser Healthcare) and the antibiotic cefazolin (330 mg/ml i.m.; West-Ward Pharmaceutical). The skin of the animal’s head covering the skull was shaved and cleaned with alcohol swabs, before the animal was placed in a stereotaxic frame (kopf). An incision was made to expose the skull and the bone tissue cleared in order to locate stereotaxic coordinates and locations for anchoring screws. Animals were implanted with unilateral microdrives containing 18-24 independently drivable tetrodes targeting the dorsal aspect of the CA1 cell layer of the hippocampus (centered at anteroposterior = -3.6 mm; mediolateral = 2.6; all coordinates derived from bregma). Each tetrode was composed of four 12 um RO 800 wires (Sandvik Kanthal HP Reid Precision Fine Tetrode Wire; Sandovik). Tetrodes were plated with non-cyanide gold solution via electrolysis in order to reduce impedance to between 180 and 220 kΩ. At the conclusion of the surgery, all tetrodes were gradually lowered ∼0.5 - ∼1.5 mm into tissue. Upon recovery from anesthesia, animals underwent postoperative care for 3 days and received doses of Buprenex and Cefazolin, as described above, two times a day (12 hour intervals). Animals were allowed to recover 1 week before behavioral re-training and tetrode lowering commenced.

### Neural Recordings

Electrophysiological recordings for this project were collected using a 96 channel OmniPlex D Neural Acquisition System (Plexon). Each channel was amplified and bandpass filtered for both single-unit activity (154 Hz to 8.8 kHz) and local field potentials (1.5 Hz to 400 kHz). Spike channels were referenced to a local electrode in the same region in order to remove both movement-related and any electrical noise. Action potentials of neurons were detected via threshold crossing and digitized at 40 kHz. Between recorded training sessions tetrodes were advanced based on visual inspection, in order to maximize neural unit yield, and allowed to settle overnight before conducting the next recording session. Individual neural units were isolated via manual offline clustering, employing Offline Sorter v3 (Plexon). Cineplex Studio (Plexon) was used for capturing behavioral tracking data via three infrared LEDs positioned atop the microdrive EIB. Cineplex Editor (Plexon) was employed to enter event markers and to verify animal positional tracking data.

### Histology

Upon completion of behavioral testing, rats were anesthetized with <5% isoflurane in oxygen. Anatomical recording sites were confirmed by creating a small lesion in the brain tissue by passing a 40 µA current until the connection was severed. Immediately after completion of the creation of electrolytic lesions, animals received an overdose injection (percutaneous) of Euthasol (Virbac AH) before being intracardially perfused with 0.9% saline followed by 10% phosphate buffered formalin (VWR). Brains were removed and placed in additional 10% formalin phosphate until sectioning into 40 µM thick sections via cryostat (CM 3050s; Leica Biosystems). Brain sections were stained with cresyl violet in order to visually confirm tetrode recording sites.

### Analysis

All analyses were performed using both built in functions and custom scripts for MATLAB version R2019a (The Mathworks) and python coding language (Python 3), utilizing the github archive https://github.com/tcnlab/maxlikespy, as well as customized code packages.

### Population level analysis

The population vector correlation analysis was conducted by first constructing average linearized firing rate vectors for all units that passed initial thresholds (minimum trial average firing rate > .01 Hz & maximum session average firing rate below 5 Hz) for each trajectory direction for each starting box location relevant to that specific recording session. The entire population recorded across the four rats used was collated in order to construct a matrix that consisted of the firing rate for each unit *j* at each linearized spatial bin *i*. For the stable box recording sessions, unit inclusion was based on even or odd trial averaged activity profiles, which was then applied to the opposite set of even or odd trial constructed firing rate vectors. This additional step was used to access the reliability of neural firing patterns of individual units included in the population-level analysis. A full trial length (combined outbound & inbound trajectory) population-based firing rate vector of either odd trial or even trial averaged firing activity was correlated (Spearman) to the opposite set of trials (even vs odd) in a bin-wise manner for the entire trial length (160 bins).

For the random box sessions, the session averaged firing rate vector for each of the separate box positions (1-6), in each direction was first constructed. Track lengths 1 (shortest) through 5 (second longest) were then correlated (Spearman) using the built in ‘corr’ function in MATLAB (Mathworks), to the corresponding linearized spatial bins of the longest track length (box position 6), at every corresponding linearized bin. This bin-wise correlation of population-based firing rate vectors resulted in a matrix of correlation values demonstrating the level of similarity of neural activity as animals traversed the linear track environment in each direction of travel for each of the six varying track lengths.

### Place field parameter estimation

The spiking properties of neurons were used to select place cells for the analysis shown in Figures 3-5. In order to evaluate to what extent the firing dynamics can be accounted for by either absolute position or position relative to the end of the track and to estimate the location and width of the of the place field in the appropriate reference frame, the presence or absence of a spike at each time *t* was quantified throughout the time the animal was navigating the track. This time series was aligned with a time series of position *x(t)* describing the linearized distance along the track in either an absolute or relative reference frame. We fit the parameters of a Gaussian spatial receptive field in each reference frame to the spike train over traversals. We found the maximum likelihood of a spike train given the model with a set of parameters ‘*Θ*. The model gives the probability of a spike at any position:

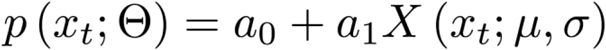

where x is position in either the absolute or relative reference frame. Factors *a*_0_ and *a*_1_ determine the contribution of each of the terms. The position term *X* is a Gaussian-shaped field defined as:

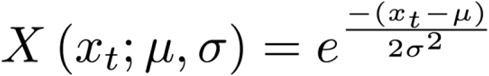

where *μ* is the spatial location of the peak of the firing field and *σ* is the standard deviation of that field.

The mean of was allowed to vary between -22.5 cm and 225 cm and the standard deviation between 0.225 cm and 202.5 cm. The bounds of the spatial parameters were chosen such that they extend beyond the dimensions of the track. In order to obtain a probability of a spike at any point in the traversal we had to ensure that the values of *p*(*x*(*t*);*Θ*) are bounded between 0 and 1. Therefore the coefficients were bounded such that a_0_+ a_1_≤ 1. The likelihood of the fit is defined as a product of these probabilities across all points in the traversal within each trial. We minimize negative log-likelihood (nLL),

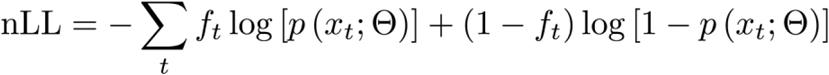

To find the best fitting models, the parameter space was iteratively searched using custom Python code implementing a global optimization routine based on SciPy’s Basin-Hopping algorithm with “TNC” as the minimization method. To avoid local minima, the parameter search procedure was repeated until a better fit was not achieved for 500 consecutive iterations.

In order to quantify whether the contribution of the spatial term was significant, the maximum log-likelihood was computed again using only the constant term. Since the models with and without spatial position are nested, the likelihood-ratio test was used to assess the probability that adding the spatial term significantly improves the fit. In order to eliminate cells with ramping or decaying firing rate, *μ* was required to be within the track boundaries.

Additionally, to eliminate cells with too few spikes, a cell was required to fire at least once per trial. Candidate cells were also fit with a time varying Gaussian model to confirm firing activity was not better explained by time relative to trial start. Cells for which the log-likelihood ratio between the best fit spatial and temporal models was above 50 were removed from the analysis. Screening for putative interneurons was achieved by eliminating cells with average firing rate greater than 5 Hz. To ensure that the model fit was robust across trials, the analysis was repeated taking even and odd trials separately. A cell was classified as either a relative position cell or absolute position cell if all the above conditions held and the likelihood ratio test was significant (p < 0.01) for even and odd trials.

## Results

### Individual CA1 units are sensitive to frames of reference

During recording sessions that employed the movable start box, individual firing fields were oriented to specific frames of reference, as has been previously reported (Gothard et al., 1996; 2001). Figure 2 shows the spiking activity of several representative units as a function of the absolute position along the linear track (top rows) or of the distance from the movable start box (bottom rows). In the outbound direction of travel (figure 2a), many “box-referenced’’ units appeared to fire at a specific distance from the start box (example units 1-3). In the outbound direction some “track referenced’’ units appear to fire at a consistent absolute location along the track (example unit 4). Qualitatively, box-referenced units span most of the journey, with track-referenced units appearing at the end of the journey as the animal approached the fixed end of the track.

**Figure 2:**
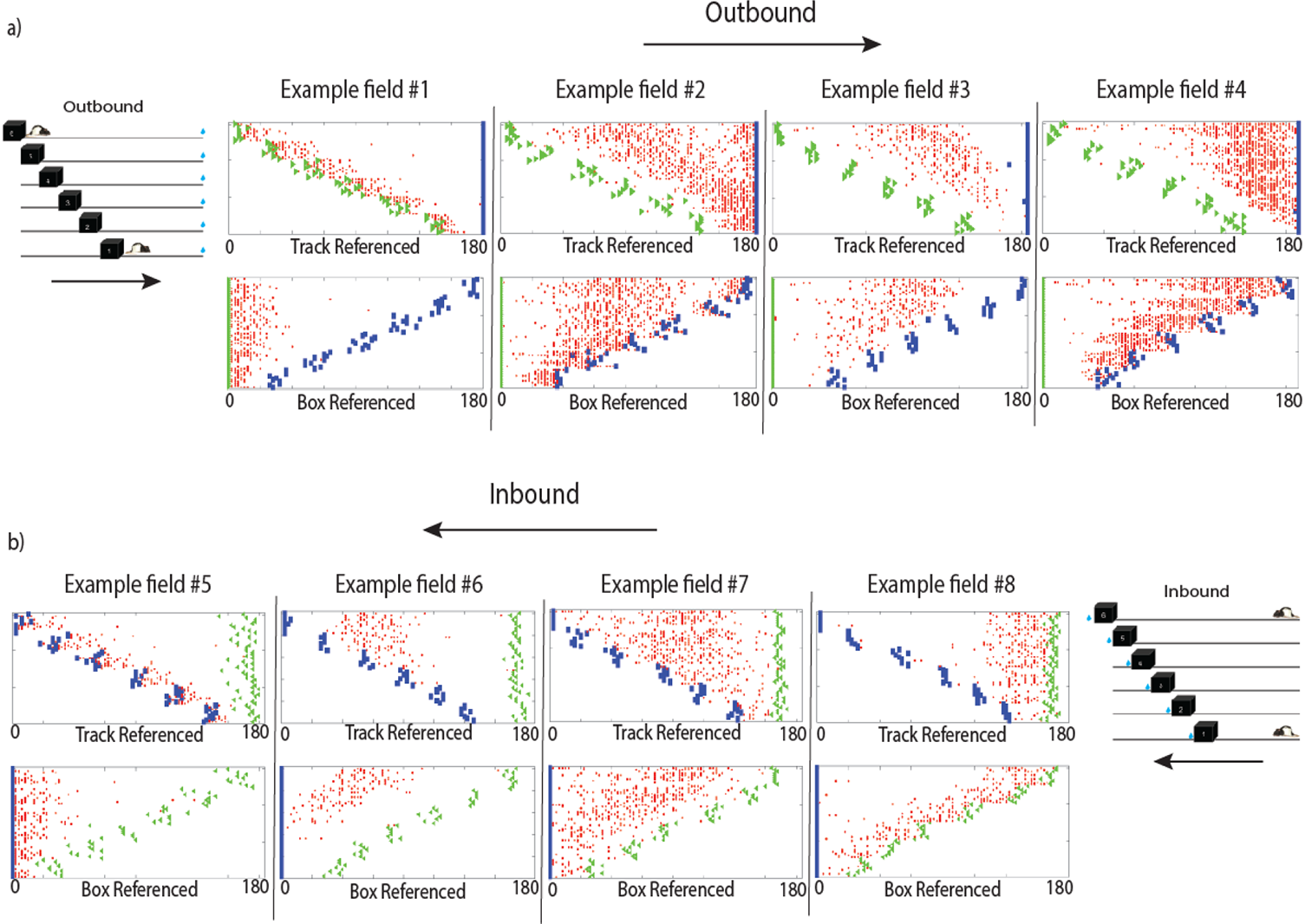
Individual hippocampal unit spiking activity during randomized start box sessions. a) Example spike rasters for individual units during the randomized start box sessions in the outbound (left to right) direction of travel, organized by starting box location (box 6: top-most block of trials; box 1 bottom-most block of trials). X-axis units denote distance in cm (full track 180 cm long). Green arrow icons denote location of trajectory start and direction of travel. Blue square icons denote water-reward markers, signaling the end of the trial. Top plots show linearized location of spikes referenced to the physical track dimensions (absolute reference frame), whereas bottom half plots show spikes aligned to the front of the edge of the moveable box (relative reference frame). Cartoon schematic of linear track environment for outbound travel during random box recording sessions. b) Example spike rasters of individual unit activity in the inbound (right to left) direction of travel. Trial organization and icons same as the above plots in fig. 2a) These rasters demonstrate many of the various activity profiles of individual putative CA1 pyramidal units observed during active track running behavior, demonstrating spiking activity in relation to several frames of reference.

In the inbound direction of travel (figure 2b), many track-referenced units appeared to fire a specific distance from the fixed starting location (example units 6-8). However, in the inbound direction there were also some box-referenced units (example unit 5). On inbound trajectories, these box-referenced fields appeared at the part of the journey as the animal entered the movable start box (bottom portion of figure 4c). Taken together, the results from the outbound and inbound journeys suggest that most place cells fire in the reference frame of the starting point of the journey, but place cells at the end of the journey fire in reference to the endpoint of the journey (Redish, et al., 2000).

### Positional coding by the hippocampal population is relative to starting location

To investigate how hippocampal neurons represented location in different reference frames during journeys along the track we examined how population vectors changed during travel along the variable length track. We compared the population vector at each position along the longest track to each position along each of the shortened track lengths (Gothard et al., 1996, 2001). By correlating the two different population vectors at each linearized spatial bin along the track we were able to determine what reference frame the hippocampal ensemble most closely reflected (figure 3). For instance, if the population code reflected the absolute position of the animal along the track, the maximum cross-correlation values of the hippocampal firing rate vectors (red line) would track closest to the dotted magenta line, which joins locations along the two journeys that have the same position in the track-based reference frame. However, if the population code reflected the animal’s position relative to the start box, the red line would track closest to the dotted green line, joining locations along the two journeys with the same position in a box-based reference frame. It is possible that the population code reflects an average of these two reference frames, in which case one would expect the red line to settle in between the magenta and green lines.

**Figure 3:**
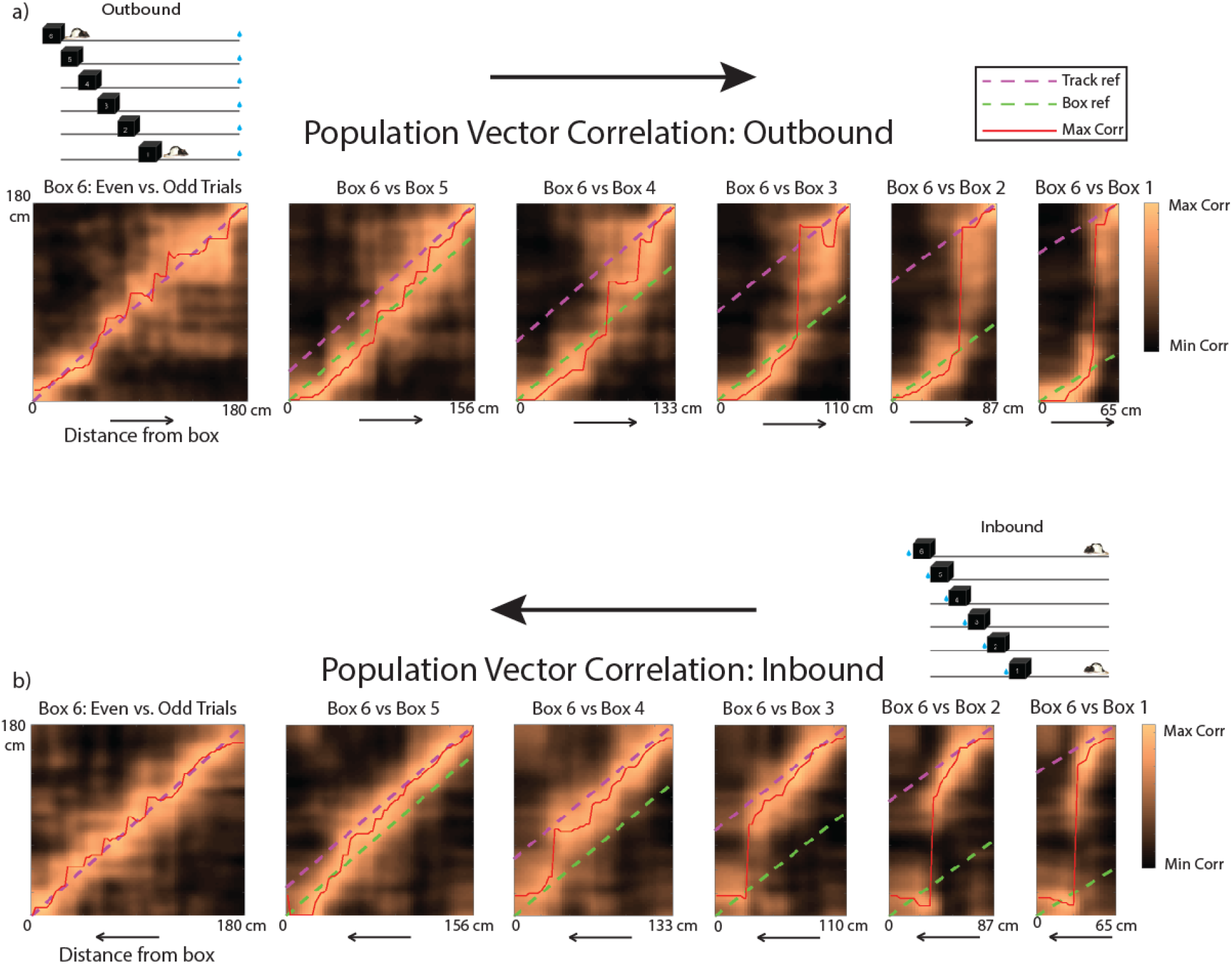
The hippocampal population represents space relative to start point. a) Outbound population vector correlations between longest track length and all subsequent shorter track lengths. Leftmost plot shows the correlation between even and odd trials for box 6 starting position, with all other subplots showing the correlation of each shorter track length, in decending order (left to right: 5-1). X-axis for all subplots is the distance from the edge of the start box to the front edge of the water reward well. Y-axis for all subplots is the total length of the longest track length (180 cm) representing the absolute linearized position. The diagonal dotted green line is the start box reference frame (relative position), whereas the dotted magenta line is equivalent to the linear track reference frame (absolute position). The red line indicates the spatial bin of best fit (highest correlation) between the two population vectors. Arrows below each subplot of a) & b) denote the direction of travel along the linear track. b) Inbound population vector correlations between longest track length and all subsequent shorter track lengths. (Same configuration and markers as above in fig. 3a). These figures demonstrate the switching of hippocampal motifs from being box referenced (i.e. relative coding) early in the trajectory, before jumping to a more track referenced (absolute coding) scheme nearest the track end in the outbound direction of travel; whereas the opposite relation of coding motif (track then box) can be observed during inbound running trajectories.

Figure 3a shows that while traveling in the outbound direction, away from the movable start box, the hippocampal representation tracked position in the box referenced frame (green dotted line: relative coding), before jumping relatively abruptly to a more track-based reference frame (magenta dotted line) as the animal approached the end of track (figure 3a). Figure 3b shows the hippocampal representation as the animals started at the fixed end of the track and returned back to the movable start box. On these inbound journeys, the hippocampal representation started out more closely aligned to the track reference frame (magenta dotted line) before transitioning to a box-centric representation (green dotted line) as the animal neared the movable start box to complete the trial. For both directions of travel across all variable track lengths, the hippocampal population code strongly represented position as a function of starting location until nearing the terminus of the running trajectory and abruptly changing to an endpoint-referenced coding motif.

### Place field allocation is biased towards the reference frame at the start of the journey

We quantified the preferred reference frame for each place cell as described in the methods. Figure 4 shows sorted place field heatmaps for the longest track length for both directions of travel. In the outbound direction of travel there were considerably more units (94.7%) with fields in the box-centric reference frame (figure 4b; bottom plot). The box-referenced fields were allocated along the entirety of the linear track length, but with more fields near the starting location. There was also a small population (5.3%) of track-referenced fields. These fields appeared to be primarily on the second half of the track, nearing the end of the outbound journey (figure 4b; top plot).

Conversely, during inbound traversals of the track a large majority (91.4%) of place fields were track-referenced (figure 4c; top). These track-referenced fields were spread across the entirety of the linear track length as well, however with more fields near the starting location. In the inbound direction of travel there was a small population of place fields that were box-referenced (8.6%), shifting their position in relation to the movable start box (figure 4e; bottom). This smaller population of hippocampal units exhibited place fields near to the movable box at the end of the journey. In both directions of travel, the majority of place fields fired in reference to the starting location of the journey (outbound: 94.7% & inbound: 91.4%).

**Figure 4:**
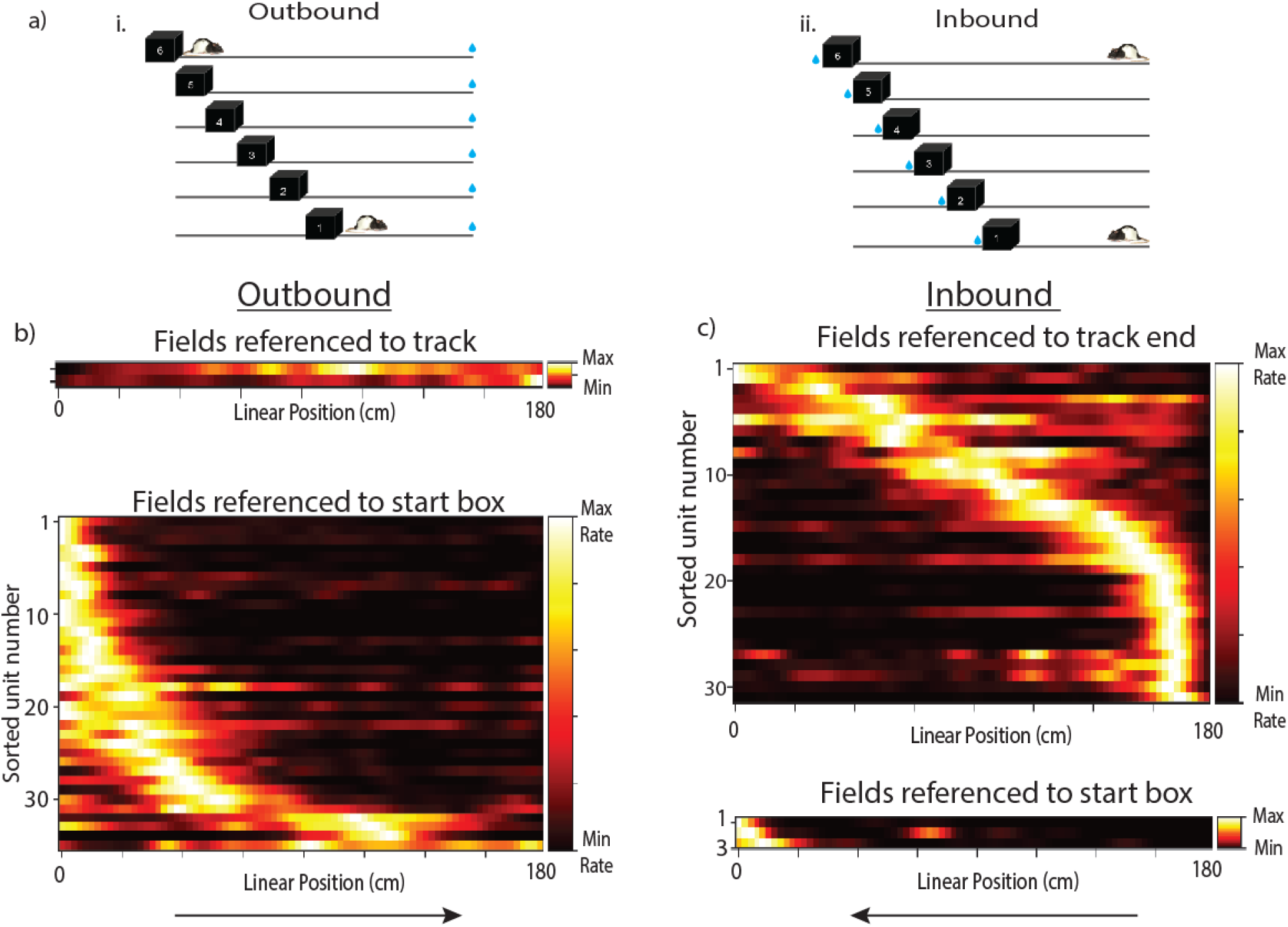
Individual hippocampal units show sensitivity to relative distance from starting location. a) Graphic of behavioral protocols employed during randomized start box recording sessions (i: outbound; ii: inbound). b) Separate heatplots for individual units that were classified as having activity fields that were either fixed to the start box (relative-based; top plot) or were absolute (track-based; bottom plot) in their firing patterns during outbound running trajectories aligned to the longest track length (box pos. 6). c) Separate heatplots for the units that were classified as either having activity fields that were track (absolute-based; top plot) or that were box-referenced (relative-based) during inbound running trajectories aligned to the longest track length (box pos. 6). Arrows at bottom of plots denotes running direction of animal during outbound (left column of plots) and inbound (right column of plots) traversal of the linear track.

### Hippocampal place fields are not uniformly distributed and express compressed activity patterns

The heatmaps of the sorted place fields depicted in figure 4 visually exhibit compression-like characteristics, in that there are more fields at the beginning of trajectories and fewer fields near the end point of travel. This can be appreciated by noting the curvature of the central ridges in Figure 4. Similarly, the widths of the fields appear to be more narrow near the beginning of a trajectory and gradually widen as the animal moves away from the starting point of the trajectory; this can be appreciated by noting the increase in the width of the central ridges as the trajectory proceeds. Figure 5 serves to quantify these visual impressions. The distribution of place field centers for units referenced to the starting position was non-uniform in both the outbound (KS-test: D=.45, p<.001, n=36; figure 5b) and inbound (KS-test: D=.29, p=.0059, n=32; figure 5c) directions of travel, as shown in the respective cumulative density plots. The two remaining populations of place fields, outbound track-referenced (figure 4c; top) and inbound box-referenced (figure 4e; bottom), did not have enough members to merit a statistical test of field distribution (5 remaining fields for each subtype).

**Figure 5:**
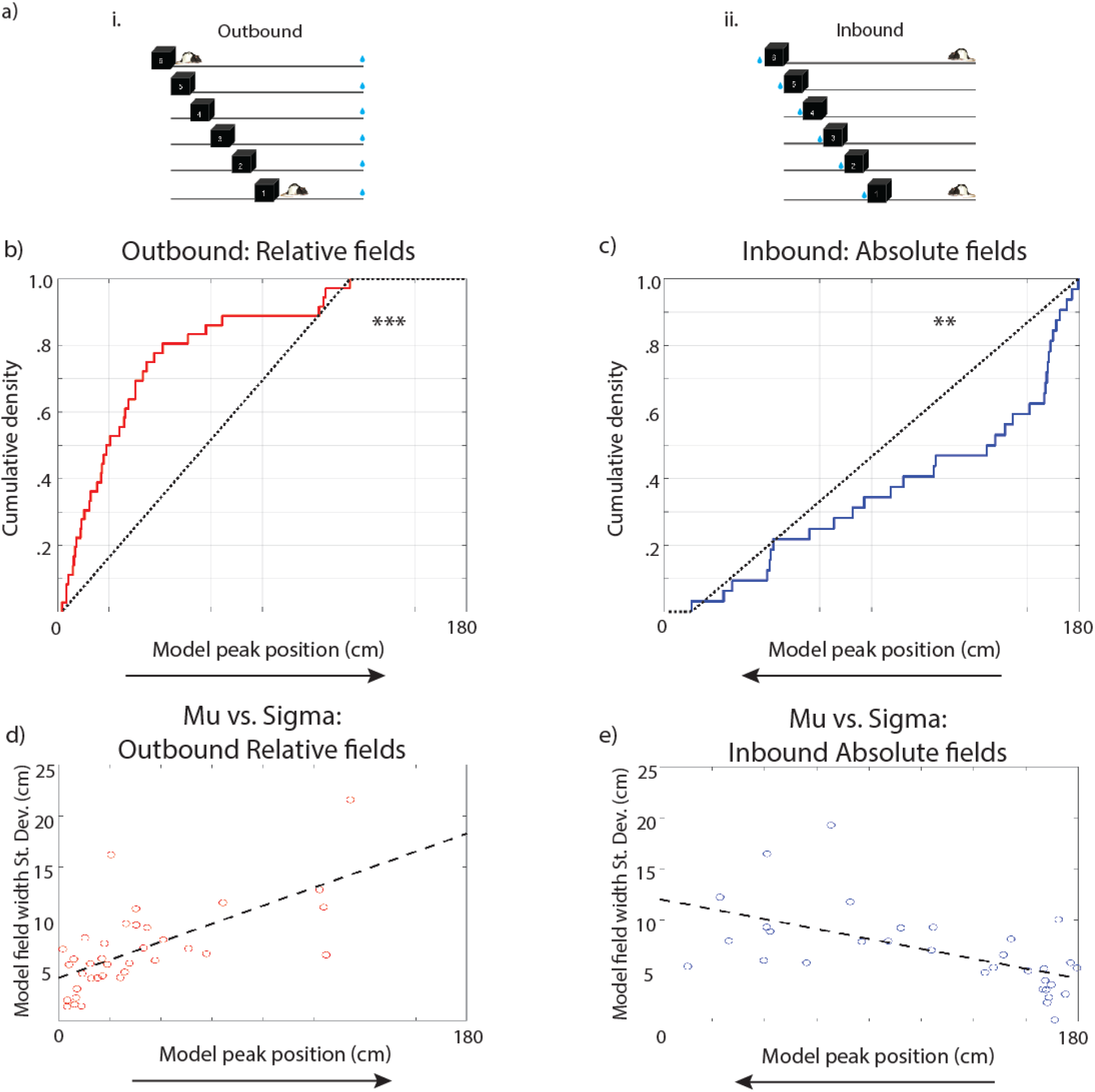
The hippocampus represents space in a compressed manner relative to starting location. a) Cartoons of behavioral protocol employed during randomized start box recording sessions for outbound (i) and inbound (ii) traversals of the track environment. b) Cumulative density plot of place field locations that were found to be relative (red line) coding during the outbound traversals of the linear track during the randomized start box recording sessions versus a bound-matched uniform distribution (black dotted line). Distribution of relative field locations in the outbound direction was found to be non-uniform (KS-test: p<.001; inset value). c) Cumulative density plot of place field locations that were found to be absolute (blue line) coding during the inbound traversals of the linear track during the randomized start box recording sessions versus a bound-matched uniform distribution (black dotted line). Distribution of absolute field locations in the inbound direction was found to be non-uniform (KS-test: p=0.006). d) Scatterplot of location of model estimated field peak (x-axis) and the St. Dev. of the field widths (y-axis) for relative coding place fields along the linear track in the inbound direction during randomized start box sessions. The black line represents the linear model that best fits these relative fields (r2= 0.432, df=34, p<.001). e) Scatterplot of location of model estimated field peak (x-axis) and the St. Dev. of the field widths (y-axis) for relative coding place fields along the linear track in the inbound direction during the return journey to the start box. The black line represents the linear model that best fits these absolute fields (r2= 0.34, df=30, p<.001). The arrow below each plot denotes the rats’ direction of travel for that respective data set.

The populations of fields referenced to the starting point were non-uniformly distributed along the track for both directions of travel. In the outbound direction of travel there is an increased density of field peaks nearest the box (figure 5b), which matches the place field heatmaps shown in the bottom portion of figure 4c. During inbound running trajectories there was a greater density of field peaks nearest the track end which was the starting location for the inbound traversals (figure 5c), which matches the place field heatmaps shown in the top portion of figure 4e. These results are consistent with a compressed representation of distance from the starting location.

Figures 5d & 5e show place field width as a function of position for the fields referenced to the starting point of each trajectory. For the outbound traversals of the track, the widths of box-referenced place fields increased with their position along the journey; a linear model showed a reliable effect of position on width (r^2^=0.43, df=34, coefficients with 95% confidence bounds: linear term= 0.17 +/- 0.07, p<.001; intercept= 4 +/- 1, p<.001; figure 5d). Similar results were observed for fields on inbound traversals of the track. Field width of the track referenced place fields increased as function of distance from the start point (r^2^=0.34, df=30; linear term - .097 +/- 0.05, p<.001; intercept= 12 +/- 3, p<.001; figure 5e). For both directions of travel, applying a quadratic term to the linear model did not improve the fit. These results demonstrate that the spatial code is represented in a compressed manner, such that positions close to the beginning of the journey are represented with more cells and more narrow place fields.

## Discussion

This study replicated and extended previous work (Gothard et al., 1996; 2001) demonstrating the sensitivity of hippocampal spatial coding to salient frames of reference. A high proportion of hippocampal place fields were anchored to the starting point of a trajectory (figures 4b & c; figure 3). Knowing the origin of the coordinate system for the hippocampal place code allows study of the properties of the place code as a function of distance from the origin. Using analytic methods closely analogous to those utilized to investigate the neural representation of time (Salz et al., 2016; Tiganj et al., 2017) we showed that individual place fields are non-uniformly distributed with higher density nearest the starting location (figures 5b & c) and that place field width increases as a function of distance from the starting location (figures 5d & e). These properties have also been observed for time cells, suggesting a common computational basis for space and time coding in the hippocampus and analogous constraints on episodic memory.

The non-uniform distribution of place field centers and increase in width are both manifestations of efficient coding of a compressed representation. Receptive fields in the visual system both become less numerous and increase in width as a function of distance from the fovea (e.g., Van Essen, Newsome, & Maunsell, 1984). If the density of receptive fields decreased without a concomitant change in the receptive field width, this would tend to insert “gaps’’ in the representation of distance, leaving regions of space that do not fall into the receptive field of any neuron. Similarly, if the density of receptive fields was constant as a function of distance, but receptive field width nonetheless increased, the result would be a highly redundant code, with the number of neurons contributing to the representation of a region of space increasing. In the visual system, it is believed that the compression of retinal space is logarithmic (Schwartz, 1977; Van Essen, et al, 1984), consistent with a Weber-Fechner scale where the neural dimension maps onto the logarithm of the physical dimension. It has been proposed that neural representations of time and space ought to share this mapping (e.g., Howard, 2017; Buszaki, 2020). Although this study did not evaluate the form of the compression, the data presented here is certainly consistent with a logarithmic compression of distance.

### Hippocampal representations of space and time

The similarity between the neural code for distance in the hippocampus evaluated in this study and the code for time evaluated in other studies suggests that the two forms of representation share a common computational basis. This is as one would expect if the hippocampus represents a general spatio-temporal context that captures relationships between the components of an event (Cohen & Eichenbaum, 1993; Eichenbaum et al., 1999; O’Keefe & Nadel, 1978). Recent years have seen a well-developed computational hypothesis for the origin of the hippocampal representation of time; some predictions of this account have been confirmed experimentally. This subsection pursues the analogous predictions for the representation of space.

It has been proposed that sequentially-activated time cells acquire their sensitivity to the time since an event from another population of cells that decay exponentially with a wide range of time constants (Shankar & Howard, 2012; Howard, et al., 2014, see also Rolls & Mills, 2019)). It has been proposed (Tiganj, Hasselmo, & Howard, 2015; Liu, Tiganj, Hasselmo, & Howard, 2019) that the variety of time constants in this population depends on intracellular Ican currents that have been observed in entorhinal neurons in slice (Egorov, et al, 2002; Fransen et al., 2006), although any other mechanism that gives rise to decaying exponential firing with a spectrum of time constants would suffice. Notably, in the last few years, populations of neurons that decay with a variety of time constants have been observed in rodent LEC (Tsao, et al., 2018) and monkey EC (Bright, Meister, et al., 2020). According to this theoretical framework, the compressed representation of time observed in the hippocampus would be inherited from a skewed distribution of time constants in the population of exponentially-decaying cells (Shankar & Howard 2013), consistent with the available data from EC.

This theoretical framework makes analogous predictions for the hippocampal representation of distance. The spatial coding of place cells is believed to be inherited from boundary vector cells (O’Keefe & Burgess, 1996, Barry et al., 2006; Lever et al., 2009) that show circumscribed receptive fields at a particular distance from an environmental boundary, much like time cells show circumscribed receptive fields a particular temporal distance from an event. The compressed representation of distance observed in this study suggests a similar compression in boundary vector cells as proposed in the original model of boundary vector cells (Barry et al., 2006; Lever et al., 2009). Border cells (Solstad, et al., 2008; Campbell, Ocko, et al., 2018) observed in MEC are the spatial analog of the population of exponentially-decaying cells in LEC (Tsao, et al. 2018). Continuing the analogy with temporal coding, this theoretical framework predicts that the distribution of space constants of border cells should show a skewed distribution analogous to the distribution of time constants observed in EC (Tsao, et al., 2018; Bright, Meister, et al., 2020).

### Reference frames and internally-generated event boundaries

Most of the place cells on the linear track had receptive fields that coded for the distance from the start point of the journey. However, a subset fired in the reference frame of the end point of the journey, with the population switching abruptly as the animal approached the end point (e.g., Figures 3, 4). The switch to the end of track reference frame may be analogous to cells in the hippocampal formation that fired a fixed distance from a reward location in virtual reality (Gauthier & Tank, 2018). In this experiment, the transition could be due to the animals switching or anchoring their “salient” frame of reference to known environmental boundaries, which could allow them to estimate relative position or distance (Hardcastle et al., 2015; Giocomo, 2016; Barry et al., 2006; Lever et al., 2009). The relatively abrupt transition from one reference frame to another along the journey (see also Redish et al, 2000) may correspond with ideas about event segmentation from the human cognitive neuroscience literature (Zacks & Tversky, 2001; Franklin, Norman, et al., 2020). Here, however, the transition was not in response to a boundary, but in *anticipation* of a boundary. Further work would be necessary to distinguish if the endpoint-referenced place cells also show a compressed representation of distance *to* the endpoint.

## Acknowledgements

This work was supported by NIMH R01MH095297, NIBIB R01EB022864, NIMH R01 MH052090, NIMH R01 MH120073 and Office of Naval Research MURI N00014-19-1-2571. This paper describes work started under the direction of the late Dr. Howard Eichenbaum. The authors gratefully acknowledge his many contributions to this paper and his inspiring leadership.

